# The aversive value of pain in human decision-making

**DOI:** 10.1101/2021.02.19.431988

**Authors:** Hocine Slimani, Pierre Rainville, Mathieu Roy

## Abstract

According to basic utilitarian principles, people should try to maximize rewards and minimize pain. Here, participants were put in a situation where monetary rewards were paired to electric shocks spanning between pain detection and tolerance thresholds. Monetary offers ranged linearly from $0 to $5 or $10 in Group1 and 2, respectively, and exponentially from $0 to $5 in Group3. The value of pain increased quadratically as a function of the anticipated pain intensity. While increasing the range of monetary offers increased the price requested to accept pain, skewing the distribution of rewards encouraged profit maximization. Participants scoring higher on harm avoidance and lower on persistence scales requested more money to accept pain. Accepting highly painful offers slowed decisions regardless of the value of the concurrent reward. Altogether, pain-related decisions are highly relative to the local range of available rewards and may be under the control of more automatic avoidance mechanisms.

## Introduction

The main function of pain is to motivate us to react to sources of actual or potential tissue damage. However, pain also marks an error in the long chain of decisions that has ultimately put us at risk of injury. From that perspective, immediate pain also teaches us to avoid future pain long before it happens. However, we may still willfully accept to withstand pain if it is associated with sufficient rewards. Indeed, according to basic utilitarian principles, rational decision-makers should try to maximize their rewards and minimize their pain (Bentham, 1996). Pain therefore has a “value” used for trading with other “goods”, such as intrinsically pleasurable activities or more secondary reinforcers like monetary incentives.

Very few studies have addressed the question of the value of pain (Talmi et al., 2009; Vlaev et al., 2009; Park et al., 2011; Winston et al., 2014), and none of them have examined its relationship with perceived pain, and how it is modulated by personality. Is the value of pain linearly associated with its intensity, or are there certain intensity ranges where pain matters more than others? To what extent do interindividual differences influence how pain is valued? These questions are clinically relevant, as the same level of pain relief could have a different value if it is felt in the high or low pain range. People who are also highly fearful of pain might assign more value to pain relief, and conversely, people who are more driven toward rewards might be more willing to accept pain for small rewards. The current study is therefore strongly embedded within the more recent developments in the Fear-Avoidance Model of chronic pain (FAM) that stresses the importance of placing pain in the context of other competing goals (Vlaeyen et al., 2016). Our long-term objective is to develop a methodological framework that would allow quantifying the value of pain beyond subjective ratings.

The present study adopted an approach inspired from behavioral economics (Padoa-Schioppa and Assad, 2006) and in which participants have to choose to accept or reject different combinations of pain and money offers. By fitting a logistic regression model to the choices, it is possible to identify indifference points, i.e. combinations of pain and money associated with a 50/50% probability of accepting/rejecting the offer. These indifference points correspond to the monetary value of a given pain level within that particular context. By tracing the curve that most parsimoniously connects all of these points, the value function of pain is obtained; i.e. the function that predicts how much pain is worth in monetary units for that individual within that particular context. In the present study, we set out to establish these value functions across a relatively large sample of healthy participants with three basic questions in mind: 1) is the relationship between pain and money linear or curvilinear? i.e. is the value of pain the same for all intensity levels, or are there some levels that have more value than others?, 2) How is the value function of pain influenced by inter-individual traits associated with harm avoidance or reward seeking?, and 3) How is the value function of pain affected by the contextual availability of monetary rewards? i.e. is pain value “absolute” or is it relative to the range of available monetary rewards, as is often observed in behavioral economics (Vlaev, 2018)? Altogether, the results of this study highlight how pain is valued in order to inform our future decisions.

## Results

### Pain sensitivity assessment

Stimulus-response curves were obtained to assess pain sensitivity. On average, the quadratic model had the lowest Akaike Information Criterion (AIC) both pre-(213) and post-test (210). The comparison of the post-valuation beta weights of this model (ß0: μ = −8.89, ß1: μ = 1.85, ß2: μ = 0.01) to the pre-valuation’s (ß0: μ = −9.85, ß1: μ = 1.99, ß2: μ = 0.01) did not show any significant difference (ß0: p = 0.098, ß1: p = 0.468, ß2: p = 0.600), indicating that pain sensitivity was stable across the experiment (d = 0.18) and hence ruling out the occurrence of significant sensitization or habituation to the electrical stimulation.

### Pain valuation assessment

Pain valuation was assessed using linear, quadratic and cubic model fits of the decision matrix. The relation between monetary rewards and the intensity of the noxious stimulus generally followed a monotonic function in all but one participant (Group 3); this data was excluded from the analysis. Overall, the AIC was lowest for the quadratic model (Group 1 = 58, Group 2 = 60, Group 3 = 57), compared to the linear (Group 1 = 60, Group 2 = 60, Group 3 = 59) and the cubic (Group 1 = 60, Group 2 = 62, Group 3 = 58) models. It therefore appears that the pain-value function is curvilinear (t = 5.04, p < 0.001). As depicted in figure 1A, the multilevel regression analysis showed that doubling the range of monetary incentives (Group 2 vs. Group 1) increased the value of pain (t = −2.08, p = 0.036, d = 0.54). However, changing the distribution monetary rewards from linear to exponential (Group 3 vs. Group 1) did not affect the overall pain valuation function (t = 1.11, p = 0.244, d = 0.29). Finally, immediately delivering rewards on every trial (Group 4 vs. Group 1) *decreased* the value of pain (t = 2.09, p = 0.018, d = 0.94). The difference between Group 1 and 2 disappeared after z-scoring the monetary rewards (Figure 1B), suggesting that the participants normalize the offers and make their decisions based on the range of the available rewards. The comparison of Group 1 and 4 could not be performed, as the multilevel model failed after normalization, probably due to the small sample size.

**Figure 1.**
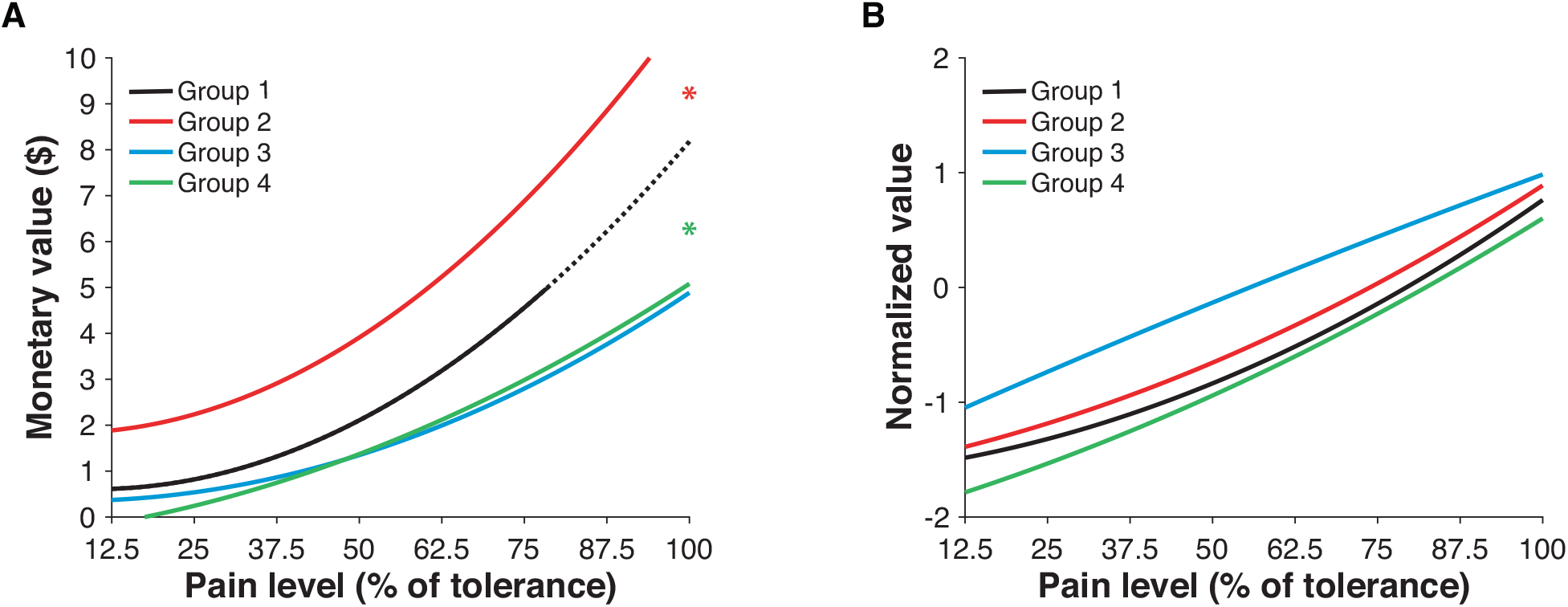
Pain value function modulation. A) The monetary value of pain increased quadratically as a function of stimulus intensity (t = 5.04, p < 0.001), with steeper increments in pain valuation when approaching pain tolerance. Group comparisons indicated that Group 2 (red) valued pain higher (t = −2.08, p = 0.361) than Group 1 (black), whereas Group 4 (green) valued it less (t = 2.09, p = 0.018). B) Normalizing the monetary incentives removed all the significant differences observed between the groups (p > 0.05).

Calculating the average acceptance rate in each group showed that the groups presented with linear reward distributions (Group 1, 2 and 4) accepted roughly half of the trials, whereas group 3 (exponential distribution of rewards) only accepted 39.6 % of the monetary incentives. However, the average gains generated by the latter were proportionally higher than those of groups 1, 2 and 4, as shown by the profitability index (Table I) calculated as follows:

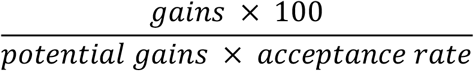

Group 3, had indeed a higher profitability index (2.03) than the three other groups (Group 1 = 1.31, d = 1.79; Group 2 = 1.34, d = 1.66; Group 4 = 1.30, d = 1.36; all p’s < 0.001). Combined with the shallower shape of group 3’s pain value function, this result suggests that the exponential distribution of rewards caused the participants to accept more easily high pain for high rewards in order to maximize their gains in the context of a relative scarcity of high rewards.

### Response times

Figure 2 illustrates the distribution of response times to the proposed pain-rewards deals. The multilevel regression analysis showed that participants were faster when offered high pain options (t = −2.52, p = 0.011). Similarly, high monetary rewards triggered faster responses (t = −5.78, p = 0.002). Responses were also faster when participants accepted offers as opposed to when they declined them (t = −5.09, p < 0.001). The interaction term between pain and decision was highly significant (t = 2.8, p = 0.004), suggesting that the participants were overall slower to accepting highly painful offers and quicker to declining them. Conversely, the decision-rewards interaction failed to predict reaction times (t = 1.71, p = 0.087). Interestingly, the pain-reward interaction predicted slower reactions when the magnitudes of both commodities were high (t = 5.16, p < 0.001), as illustrated by the top-right corner of figure 2. Surprisingly, choice difficulty, as indexed by the proximity of the trial to each participant’s indifference point, did not affect response times (t = −1.35, p = 0.166).

**Figure 2.**
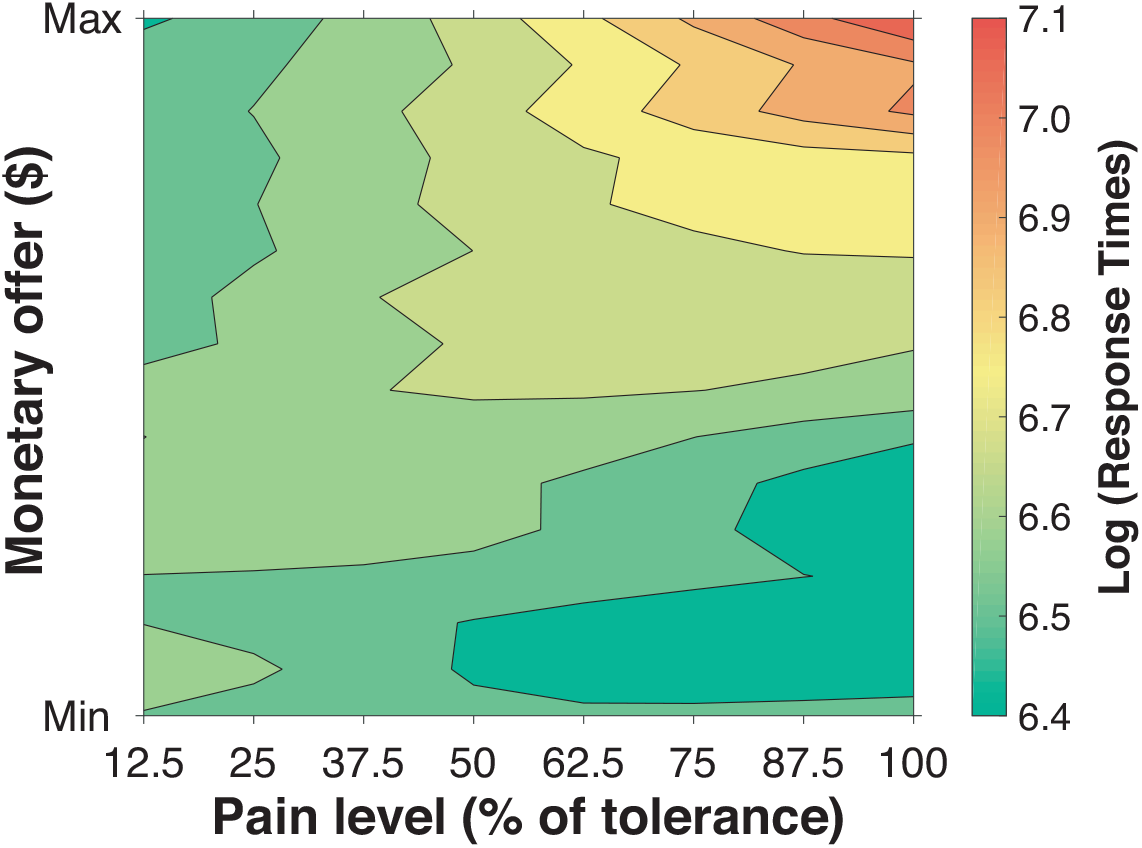
Pain valuation response times. Logarithmic average response times during the pain valuation task, with the green-to-red scale indicating a faster-to-slower decision-making, respectively. The table shows that the choice difficulty index did not significantly affect response times (t = 1.35, p = 0.166). On the other hand, we found that response times were faster when participants accepted offers as opposed to when they declined them (t = −5.09, p < 0.001). Response times were also faster when participants were offered high pain options (t = −2.52, p = 0.011), as well as high monetary rewards (t = −5.78, p = 0.002). Interestingly, the interaction term between pain and decision was highly significant (t = 2.8, p = 0.004), suggesting that it took longer for the participants to accept a highly painful offer than to decline it, regardless of the monetary offer. Finally, when looking at the interaction between the choice and the magnitude of the monetary offer, we found a trend towards faster reaction times when accepting offers of higher rewards (t = 1.71, p = 0.089) as opposed to rejecting them.

### Impact of personality traits

The PCA of the 11 questionnaires subscales generated 11 orthogonalized principal components; the 5 first explaining 92.5 % of the variance. The first component, explaining 43.4 % of the variance was mainly driven by the Trait and Character Inventory (TCI)_harm-avoidance_ (r = 0.755) and the TCI_persistence_ (r = – 0.504) subscales, suggesting that it tracked harm avoidance in a way that interferes with the capacity to persist in the face of challenges. Component 2 accounted for 19.6 % of the variance and tracked the TCI_novelty-seeking_ (r = 0.739) and the TCI_persistence_ (r = – 0.596). Component 3 (12.0 %) and 4 (9.9%) tracked the State and Trait Anxiety Inventory (STAI)_state_ at r = 0.523 and r = 0.714, respectively. Component 5 (7.7%) tracked Fear of Pain (FOP)_severe_ (r = 0.679) and FOP_medical_ (r = 0.587). Out of the 5 selected principal components, only the first one related to harm avoidance and persistence influenced decisions. Individuals scoring higher on the harm avoidance and low on the persistence dimensions were more likely to reject the offer (t = 2.04, p = 0.016), suggesting that pain has a higher value (Figure 3A). In addition to personality traits, we also tested the influence of all of our sociodemographic variables, including socioeconomic status, and found that none of them influenced participants’ valuation function (all p’s > 0.05). We also examined the influence of inter-individual differences in current intensity and pain rating at the tolerance threshold (Figure 3B), and found that the individually calibrated pain tolerance level, but not current intensity, influenced the pain-valuation function (effects of pain rating at tolerance: t = 2.04, p = 0.005; effects of current intensity at tolerance: t = 0.88, p = 0.482). These results indicate that participants who received higher pain offers as a result of having a higher pain rating at tolerance during the calibration procedure also tended to value those higher pain offers more, i.e. required more money to accept them despite the fact that they were a constant fraction of the tolerance threshold across individuals.

**Figure 3.**
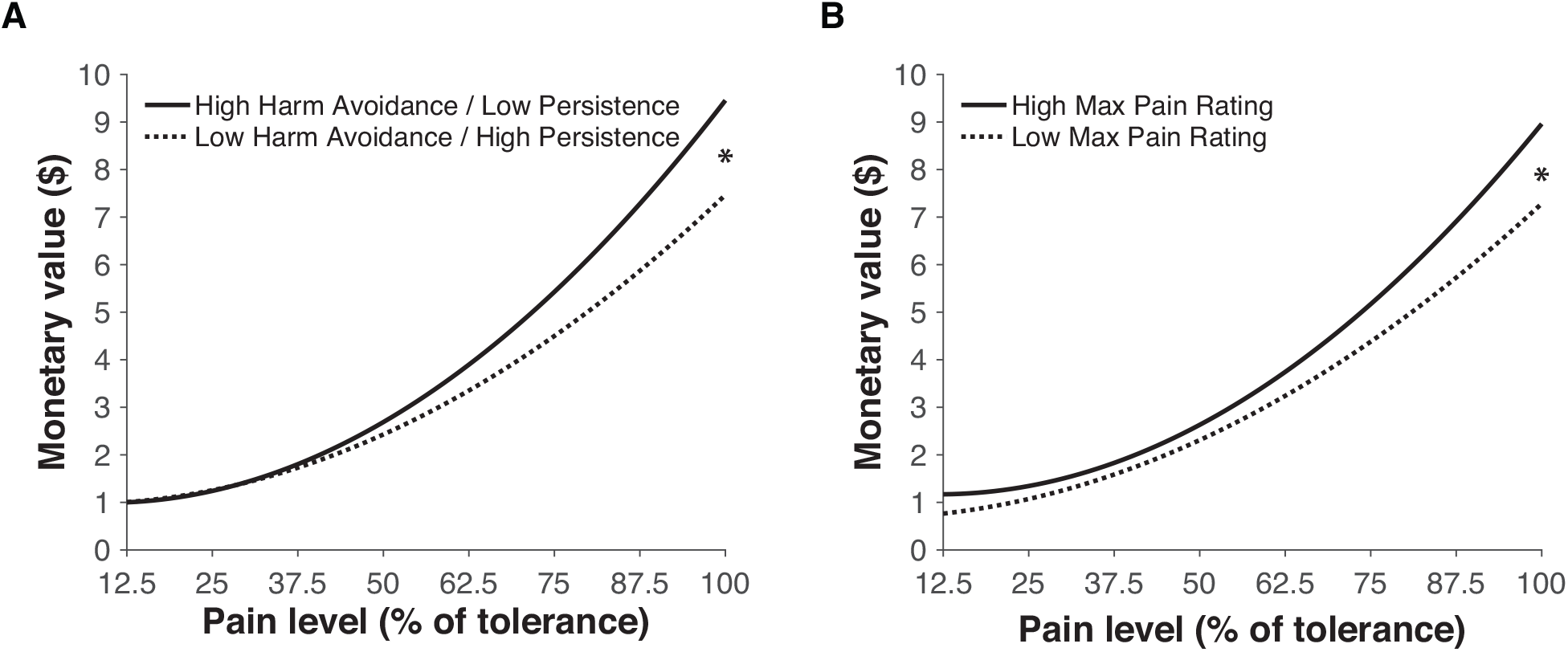
Interindividual variability of the pain value function. A) The dotted and full lines respectively represent the pain value function of the 25^th^ and 75^th^ percentiles of the first principal component of the PCA. This component was mainly driven by the TCI_harm-avoidance_ (r = 0.755) and the TCI_persistence_ (r = – 0.504) subscales, indicating that people who score higher on the “Harm Avoidance” and lower on the “Persistence” subscales of the Trait and Character Inventory attribute a higher monetary value to pain (t = 2.04, p = 0.016). B) The dotted and full lines respectively represent the pain value function of the 25^th^ and 75^th^ percentiles of the pain ratings at tolerance level. The higher people rate their pain at tolerance, the higher they value pain (t = 2.04, p = 0.005).

## Discussion

According to Bentham’s famous formula, pain and pleasure are the “two sovereign masters” under which nature has placed mankind. While several studies have investigated how rewards drive decisions (O’Doherty et al., 2017), much less is known about pain. In this study, we investigated how the value of pain is compared against that of monetary rewards during decision-making. Results show that pain-value function is quadratic, and that it is strongly influenced by context and personality. Moreover, accepting more painful offers slows decisions regardless of the value of competing rewards, suggesting that pain-related decisions may not be entirely under the control of instrumental decision-making systems. Finally, personality traits related to harm avoidance and the calibrated tolerance level also influenced participants’ willingness to accept pain in the perspective of future rewards. Altogether, the results of the current study open the way for a larger research program aimed at understanding pain-related decision-making by showing that pain is largely similar to other economic goods, with some potentially interesting exceptions that might reveal new influences not detected by classical paradigms.

This is to our knowledge the first study to outline the pain value function, i.e. to delineate how much each unit of pain is worth in terms of money, and by extension any type of good that can be exchanged for money. The pain value function is slightly curvilinear: it increases faster with increasing levels of pain intensity. This is interesting because this would suggest that the same degree of pain relief (e.g. 10/100) has more value when it is in the higher vs. lower pain range (e.g. from 90/100 to 80/100 vs. 20/100 to 10/100). Quadratic effects were small, but robust across our 4 experimental groups. We were, however, limited in the range of painful intensities that we could offer to our participants. Stimuli could not exceed participants’ tolerance limit, defined *a priori* during the calibration phase of the experiment. This tolerance limit corresponds to how much pain participants would be willing to take in the general context of a scientific experiment without additional monetary rewards. Arguably, participants could have accepted more pain provided they were given sufficient incentives, and we therefore predict that quadratic effects would have been more pronounced if we could have sampled offers above their initial tolerance limit. This suggests that the value of pain could rapidly reach very high levels as we venture past the tolerance limit, although this may be ethically impossible to confirm because it would entail making offers above participants’ natural tolerance level.

The value of pain also seemed to be highly contextual. Indeed, participants appeared to adjust their choices as a function of the mean and standard deviation of monetary offers. This caused them to request more money for pain when the range of available rewards was high, and conversely less money for pain when the range of available rewards was low. This is in line with the notion that people have comparative representations of utility, even for subjective aversive experiences (Ariely et al., 2003; Vlaev et al., 2009; Vlaev et al., 2011; Vlaev, 2018). In the same way that our perceptual systems need to re-scale sensory inputs in order to map the extremely wide range of physical stimulus intensities that we are exposed to, our valuation system also needs to re-scale the value of the goods that we are currently choosing from (Khaw et al., 2017). For instance, we may spend considerable energy choosing what to order for dinner, despite the decision being far from a matter of life and death. Therefore, like other goods, the value of pain seems to be highly menu-dependent (Padoa-Schioppa, 2009). This contextual re-scaling of value seems to be deeply rooted in the scaling properties of our neurophysiological decision-making systems, which will adjust their firing rates as a function of the range of available rewards (Tremblay and Schultz, 1999; Nieuwenhuis et al., 2005; Tobler et al., 2005; Seymour and McClure, 2008; Vlaev et al., 2011). This phenomenon, called “range adaptation”, ensures that the full activity range is always available to represent the range of values offered in the current context and hence provides an efficient representation of available rewards (Rustichini et al., 2017).

The results further show that when high rewards are scarce, participants maximize their gains by accepting more pain to obtain the rare high rewards, just as consumers may be tricked in purchasing certain goods that “stand out” by lowering the relative value of available alternatives (Tversky and Kahneman, 1974). This suggests that decisions about pain could therefore be easily manipulated by the same marketing strategies proven to be effective with for other sorts of goods. For instance, decoys (Huber et al., 1982) and other types of anchoring effects (Sherif et al., 1958) could be harnessed to render pain more acceptable in clinical settings, such as in exposure-based therapy for pain (McCracken and Vowles, 2006; Vlaeyen et al., 2012), or physical exercise therapy (Geneen et al., 2017; Qaseem et al., 2017). The current findings could therefore continue to pave the way for a larger research program aimed at understanding how people make decisions about pain (Vlaev et al., 2009; Vlaev et al., 2014; Vlaeyen et al., 2016; Vlaev, 2018). As mentioned earlier, the impact of pain on behavior resonates far beyond the immediate pain experience, and most of the maladaptive coping mechanisms observed in patients with chronic pain may derive from decisions about anticipated, i.e. *imagined*, pain. It therefore seems primordial to understand how pain’s value is represented and compared against that of other economic goods. While, the importance of competing sources of motivations had already been stressed out in recent reformulations of the fear-avoidance model of chronic pain (Vlaeyen et al., 2016), the present results show that the local range of available rewards can have a strong impact on pain-related decisions, and that it should therefore be relatively easy to influence pain behavior by manipulating the local context of competing rewards.

Perhaps the most impressive expression of these contextual effects was the behavior of group 4 (immediate total rewards), who could have gained an additional 117$ in average (see Table 1) by simply accepting pain offers that were often well below their tolerance threshold. By comparison, participants in group 1 (delayed fraction of rewards) accepted about the same amount of pain for a much smaller fraction of rewards. Contextual manipulation of competing rewards therefore seems to make a real and significant difference on participants’ decision to accept or avoid pain. It is likely that manipulating the range of pain offers would have led to similar scaling effects of pain value. However, the range of intensities that we can administer in experimental settings is limited by ethical constraints, and the present study therefore opted to start by mapping the largest possible portion of pain-value function. Still, a systematic manipulation of the range of pain offers could easily be implemented in a subsequent study. Based on the present findings, we would predict that higher ranges of pain offers would lead participants to accept higher pain intensities for less money.

**Table 1.** Profitability index.

| | Potential Gain<br>(\$) | Average Gain<br>(\$) | Acceptance Rate<br>(%) | Profitability<br>Index |
| --- | --- | --- | --- | --- |
| **Group 1** | 320.00 | 202.05 | 49.6 | 1.31 |
| **Group 2** | 640.00 | 412.28 | 49.6 | 1.34 |
| **Group 3** | 141.68 | 102.43 | 39.6 | **2.03** |
| **Group 4** | 320.00 | 203.00 | 49.3 | 1.30 |

Digits in bold illustrate significant differences from Group 1 ( $p < 0.001$ ).

Overall the current findings are consistent with the fear avoidance model (FAM) of chronic pain. Indeed, decisions about pain were strongly influenced by expected pain and rewards, i.e. the offers themselves, as well as by the learned, or inferred, range of potential pain and rewards. Experienced levels of pain or monetary rewards may only have an impact on behavior through their influence on expectations – *inferences* – about future pain and rewards. This leaves a lot of room for the influence of psychological factors beyond immediate pain sensitivity, which coincides with the main tenet of the FAM: what drives the maladaptive avoidance of pain is the fear associated with expectations about future pain (Vlaeyen et al., 2016). Here, the effects of expected pain were not always completely relative to the range of pain offers. Indeed, participants who received a higher range of pain offers due to higher tolerance thresholds also requested slightly more money to accept these higher pain offers. This means that manipulating the range of pain offers might not lead to a complete re-scaling of pain values, as was observed with monetary rewards. Still, this interesting hypothesis should be further confirmed by studies designed to directly compare the impact of range manipulations across pain and monetary offers.

Potential differences in decision-making between pain and money are also very interesting from the standpoint of behavioral control systems. By definition, decisions have to be driven by an instrumental decision-making system that learns to maximize rewards and minimize pain. However, pain is also evolutionary designed to prompt automatic avoidance of the actions that just caused it, regardless of competing rewards. Indeed, decisions taken to avoid excruciating pain generally feel coerced, i.e. not willfully taken by our instrumental decision-making system. This property of pain, along with other primarily aversive stimuli, has led certain researchers to propose that decision-making in aversion may be more prone to the disruptive influence of an additional Pavlovian system (Dayan and Seymour, 2009). The idea of an accelerating pain value function that eventually reaches an asymptote is consistent with this hypothesis of a more stable referent tied to the inherent biological meaning of pain. Moreover, we observed that response times did not perfectly matched decision difficulty but were rather driven by the interaction of pain level and decision: it takes longer to accept high levels of pain. This suggests that participants might hesitate more before self-administering highly painful shocks, regardless of the whether or not the deal is good from an instrumental standpoint. Fear of pain might therefore produce some decision-theoretical anomalies when making decisions about pain whereby there is an innate tendency to avoid pain regardless of the rewards that may be associated with it. This could also explain the impact of personality traits on decisions about pain: participants with high level of harm avoidance and low levels of persistence required more money to accept the same levels of pain. This suggests that the capacity to resist the dread of pain might be important for obtaining desired delayed rewards in spite of immediately threatening pain. However, this hypothesis would still need to be tested in future studies systematically manipulating the nature (appetitive vs. aversive) and temporal discounting (immediate vs. delayed) of outcomes.

Altogether, the results of our study indicate that pain indeed has a value that can be traded for other goods. Moreover, this value appears to be highly relative to range and distribution of available rewards. This context-dependency presents many opportunities for interventions aimed at reducing the occupational limitations imposed by pain. That being said, there also seems to be certain limits to our capacity to willfully accept pain. By design, pain can also be imperative: it dictates avoidance (Klein, 2015). While the possibility that decisions about pain are also influenced by a Pavlovian system still requires further demonstration, this could also lead the way to more tailored interventions for influencing decisions about pain. Overall, the current study therefore provides some of the theoretical framework, and basic methodology, for a vaster research program aimed at understanding how pain influences decisions in the face of competing rewards.

## Materials and Methods

### Participants

Ninety healthy participants (45 F) aged between 18 and 40 years were recruited through advertisements posted in the campuses of McGill University and Université de Montréal. Exclusion criteria included any current neurological, psychiatric, or pain-related disorders, as well as regular consumption of psychotropic or analgesic medication. The protocol was approved by the ethics committee of the Centre de Recherche de l’Institut Universitaire de Gériatrie de Montréal (CRIUGM), where the study was performed (CER VN 16-17-12). Participants were given a monetary compensation of $45 for their participation.

### Procedure

The participants were enrolled in a single testing session of approximately 3 hours at the pain laboratory of the CRIUGM. The study began with participants completing a battery of questionnaires assessing different personality traits related to pain and reward processing. Thereafter, they underwent a familiarization procedure where they were shown the electrical stimulator, explained the different steps of the experiment and given sample stimulations. Then, participants underwent a calibration procedure where they rated the pain produced by electrical stimulations of various intensities so as to determine the intensities of the stimuli to be used for the decision-making experiment. Next, participants performed the main decision-making task where they repeatedly had to accept or decline pairs of painful shocks and monetary rewards. Finally, we reassessed their pain sensitivity in order to monitor any possible changes in pain sensitivity.

### Questionnaires

The participants completed a battery of online questionnaires including the Sociodemographic Questionnaire (SDQ), the Pain Catastrophizing Scale (PCS, Sullivan et al. (1995)), the Fear of Pain Questionnaire (FOP, McNeil and Rainwater (1998)) the State-Trait Anxiety Inventory (STAI, Spielberger (1983)), and the Novelty Seeking, Harm Avoidance and Persistence subscales of the Temperament and Character Inventory (TCI, Cloninger et al. (1987)). The State Anxiety subscale of the STAI (STAI_state_) was administered between the pain calibration procedure and the pain valuation task in order to measure the impact of the electric shocks on the participants’ anxiety levels. The PCS was also administered during that time as a situational assessment of catastrophic reactions to the painful electric shocks used in the experiment (Campbell et al., 2010; Leung, 2012). All other questionnaires were answered before being exposed to the electrical stimulations.

### Calibration procedure and pain ratings

The calibration procedure started with a familiarization phase where the experimenter began by introducing the apparatus and explaining the experimental procedure. The participant then sat comfortably on a dentist chair. Stimulation electrodes (1 cm^2^, distance of 1 cm between electrodes) were placed on the degreased skin of the retromalleolar path of the right sural nerve. Nociceptive stimuli consisted of 30 ms transcutaneous electrical shocks (trains of 10 1-millisecond pulses at 333 Hz) delivered with an isolated DS7A constant current stimulator (Digitimer Ltd, Welwyn Garden City, United Kingdom) triggered by a S48 train generator (Grass Medical Instruments, Quincy, MA) and controlled by a computer running E-Prime2 Professional (Psychology Software Tools, Sharpsburg, PA). A classical staircase method was used to obtain estimates of pain detection and tolerance thresholds. More specifically, the experimenter started with a 1-mA stimulus and gradually increased the intensity by 2 mA. A visual analog scale (VAS) was used to indicate the pain level elicited by each electrical stimulation (0: no pain to 100: extremely painful). The VAS consisted in a graduated horizontal bar shown on the computer screen with a cursor moved using a computer mouse. The participants were instructed that the intensity would slowly increase after each trial until reaching their tolerance level.

### Pain sensitivity assessment

The intensity corresponding to each participant’s pain tolerance level was divided by 8 to determine 8 evenly distributed shock intensities between the pain detection and tolerance thresholds. The tolerance rating is here defined as an individually-adjusted number referring to the VAS rating reported by the participant when they asked the experimenter not increase the intensity anymore in the calibration phase. This was done to account for individual differences in the VAS-tolerance matching point. Electric shocks were randomized in blocks that included all the possible intensities using a random list generator (https://www.random.org/lists/). This avoided the presentation of the same intensity level more than twice in a row. The participants were submitted to 4 blocks of stimuli for a total of 32 trials. After each electrical stimulation, participants rated their pain on the VAS. Next, lasso regression with a 32-fold cross-validation procedure was used to determine the parameters of the power function that best describes the participant’s pain sensitivity (Matlab Statistical Toolbox (Matlab R2018a, The MathWorks Inc). This personalized stimulus-response function was then used to select 8 shock intensities corresponding to 8 levels of pain ranging from 12.5 to 100 % of the participant’s pain tolerance (Figure 4). Finally, this procedure was repeated after the pain valuation task, in order to monitor possible changes in the participant’s pain sensitivity between the beginning and the end of the experiment.

**Figure 4.**
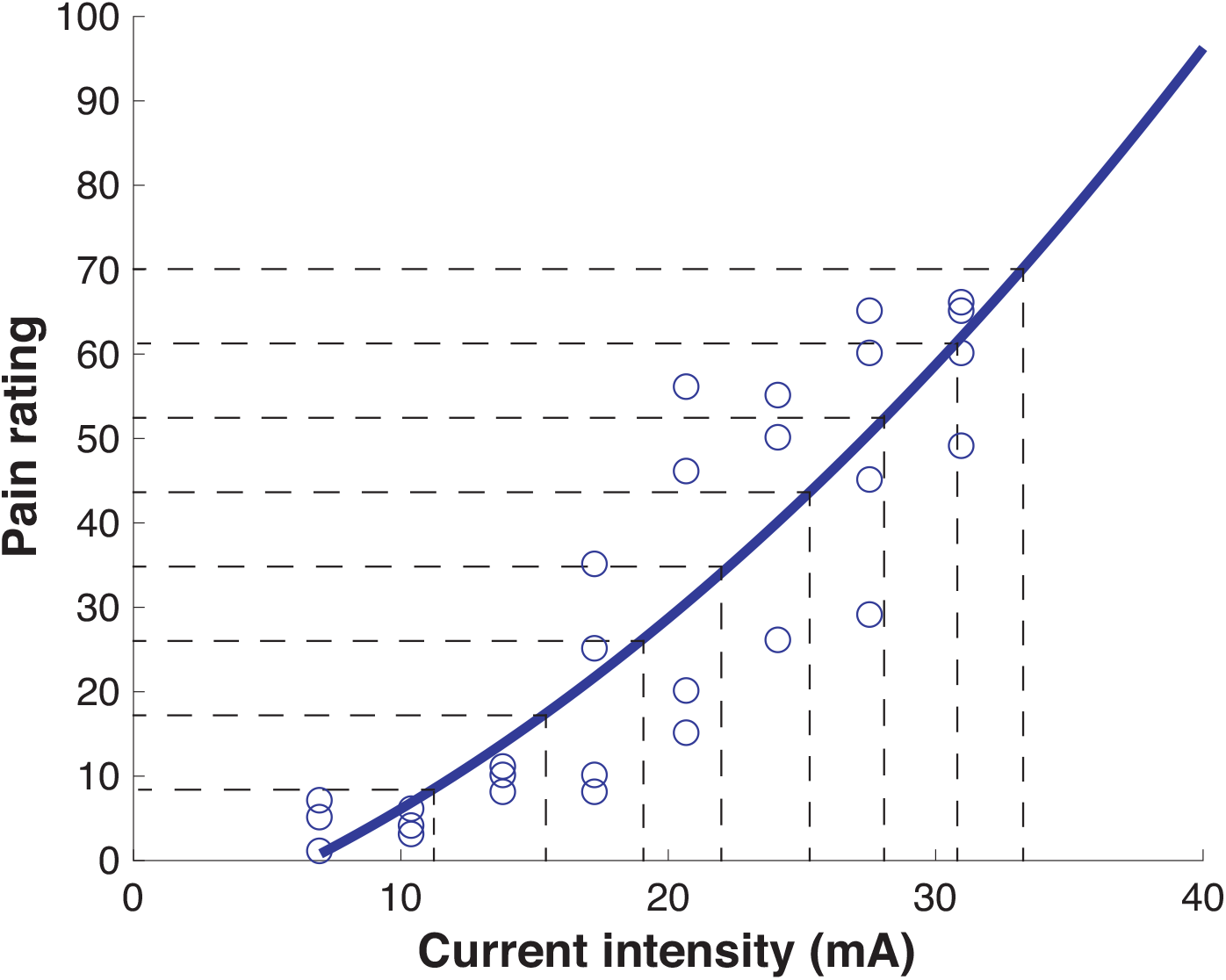
Example of a pain sensitivity function. Participants experienced each of 8 different levels of electric shocks 4 times in a pseudo-randomized order. The data was fitted after a model selection using a lasso regression analysis. The shock intensities corresponding to 12.5 to 100% of tolerance (here = 70/100) were determined based on the sensitivity function generated by better fitting the general linear model.

### Decision-making task between pain and money

In the experimental task, participants had to accept or decline painful shocks that were paired with monetary rewards. The painful stimuli consisted of 8 equally spaced intensity levels ranging between 12.5 to 100 % of the participants’ pain tolerance level, as determined in the pain sensitivity assessment procedure. These 8 levels of pain were paired were 16 levels of monetary rewards for a total 128 trials. In order to assess how choices are influenced by the range of available rewards, we varied the maximum and the distribution of monetary rewards across the 3 groups. In Group 1 and 3, offers varied between $0 and $5, with a linear or exponential distribution of rewards, respectively. In Group 2, the offers increased linearly from $0 to $10 (Figure 5A). As often performed in behavioral economics task (Chib et al., 2009; Park et al., 2011), participants were told that they would receive the monetary rewards at the end of the task based on a random selection of a limited set of trials. If participants had accepted the pain offers on the selected trials their corresponding amount of money would be compiled and added to the $45 baseline compensation. This procedure prompts participants to treat every trial as if it was one of the selected trials. Moreover, participants were unaware of the number of trials the task required in order to avoid the use of global strategies that would hinder their ability to make punctual decisions, as instructed. They also had to learn the range of pain stimuli and monetary rewards, as they were not informed *a priori* of these variables’ minimum and maximum. Finally, in order to check the impact of that procedure on participants’ decisions, we tested 6 participants (Group 4) who directly received monetary rewards ($0 to $5, linearly distributed as in group 1) in an opaque jar placed in front of them. Since all compensations were given upon acceptance of a deal, the potential gain per participant for this experimental manipulation was of $320. The sample size was hence limited by budgetary restrictions. Given the low sample size, data from group 4 was not used in all the analyses, and mainly served as a manipulation check against potential biases related to the delayed administration of rewards in groups 1-3.

**Figure 5.**
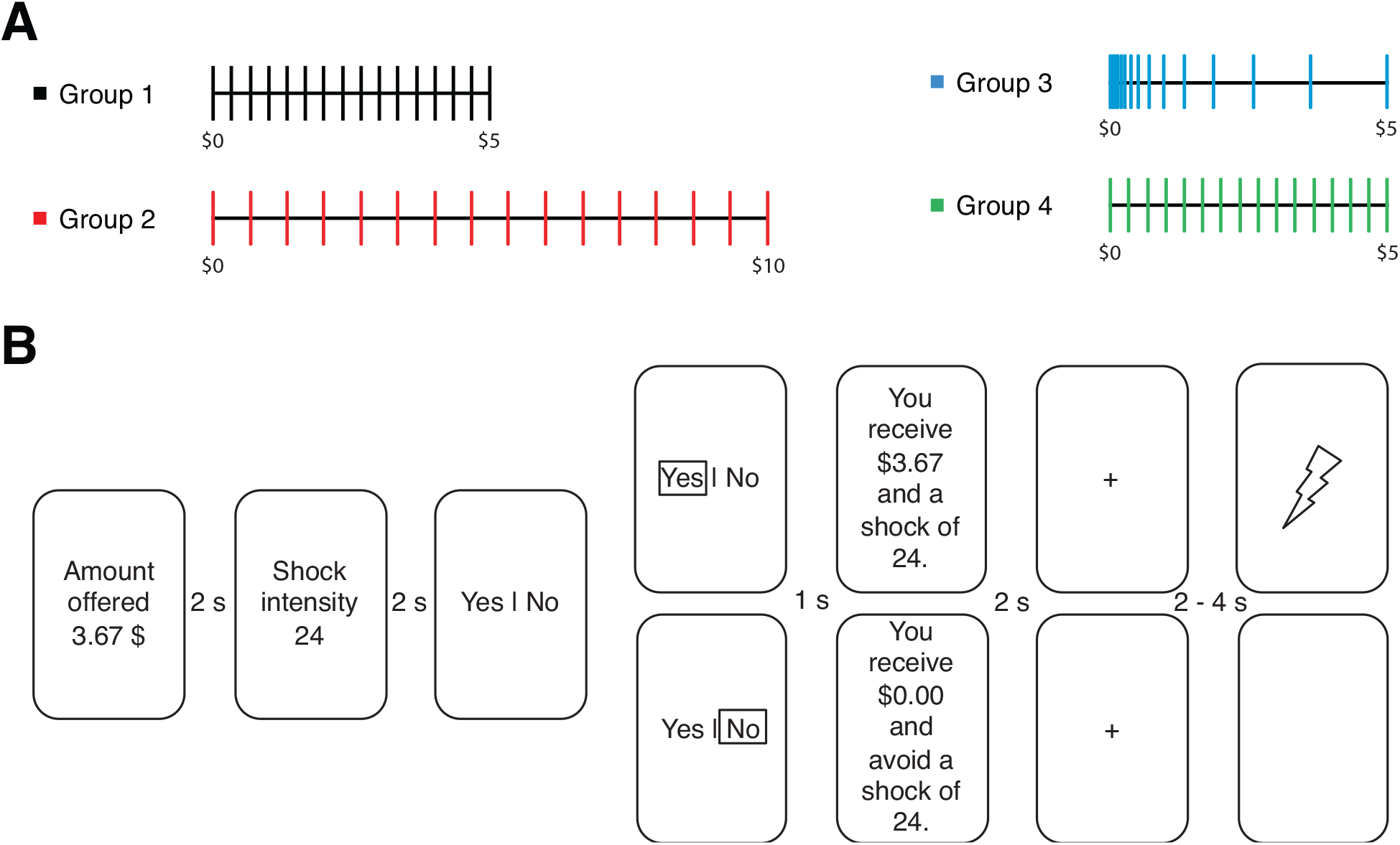
Pain valuation assessment. A) A money-first trial where participants were presented with a monetary offer, followed by a pain offer (inversely for a pain-first trial). Upon acceptance of the deal, they would receive an electric shock of the announced pain intensity and the monetary reward would be stored in a bank to pick from randomly (Groups 1,2 and 3) or received right away in an opaque jar. Inversely, declining the offer lead to avoiding the shock and missing the opportunity of a reward. B) The 16 monetary offers ranged linearly from 0 to $5 in Group 1 and 4 or $10 in group 2. They increased exponentially from 0 to $5 in Group 3.

Each trial began by the presentation of one of the two offers, pain or money, followed by the second offer (50% each; order counterbalanced). This procedure was used in prevision of a future brain imaging study that would examine the brain’s response to monetary and pain offers separately. For example, in a money-first trial (Figure 5B), participants were first presented with a monetary offer for two seconds. Then, following a 2-s interval, they received the pain offer, which was displayed on screen for another 2-s. Finally, after another 2-s interval, they were then prompted to accept or decline the offer. Response times were recorded. When accepting the offer (top panel in Figure 5B), participants received the announced electric shock 2-to 4-s later (jittered interval; uniform distribution between 2.000s and 4.000s). Conversely, no electric shock was administered following a declined offer, but participants had to wait the same amount of time before moving to the next trial.

### Statistical analyses

#### Effect size

Statistical analyses, including Cohen’s d (d) calculations for effect size, were done in MATLAB (2018a) using custom code available at https://canlab.github.io/.

#### Pain sensitivity

The pain sensitivity assessment was performed at the beginning and the end of the experiment to monitor any changes in perceived pain over time. Paired Student T-Tests were performed on the function’s ß-weights generated by the lasso regression analysis to compare the pain sensitivity pre- and post-valuation.

### Pain value function

Decisions were first arranged as an 8 (pain offers) by 16 (monetary offers) matrix, which was then smoothed using a smoothing kernel. Then, a decision matrix was fitted with a logistic regression model to predict choices from the offered levels of pain and money. Trial effects were controlled for by adding the trial number, the log(trial number) and the reverse order log(trial number) as a covariates. In order to account for potential curvatures in pain’s value function, linear, quadratic and cubic models were tested and compared at the group level using Akaike Information Criterion (AIC). The indecision data points corresponding to a 50% likelihood of accepting an offer from the fitted decision matrix provided a pain value function for each individual (Figure 6). Finally, a multilevel regression analysis with “Group” as the independent variable was computed to evaluate the impact of the range/distribution of monetary offers on the pain value function (dependent variable).

**Figure 6.**
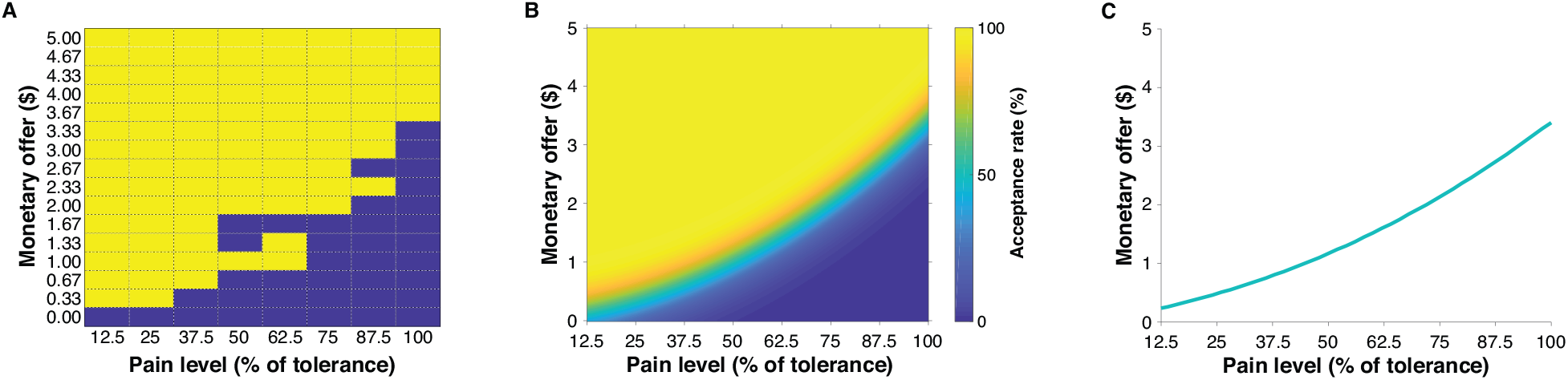
Pain value function. A) Raw data from one participant illustrating the acceptance (yellow) and rejection (blue) of different combinations of monetary offers and electric shock intensities. B) Quadratic modelling of the participant’s decision matrix. The color code represents the probability of accepting an offer. C) Extraction of the indecision points of the fitted matrix, namely the pain value function.

### Response times

The response times matrix was arranged similarly to the decision matrix: 8 pain levels by 16 monetary rewards. The predictive value of the trial number, the log(trial number) and the reverse order log(trial number) on the response times was assessed with a general linear model. By keeping the model residuals, we rid the response times matrix of order effects. The response times matrix was then smoothed using a smoothing kernel. Finally, a multilevel regression analyses was computed to evaluate the impact of the following variables on the reaction speed (1/response times): pain intensity, monetary reward and decision (accept vs reject the offer), as well as the interaction between pain intensity and monetary reward, pain intensity and decision, and monetary reward and decision.

### Questionnaire data

Participants were screened for personality traits related to pain processing and reward seeking. Data from 9 participants was excluded because of they incorrectly responded to the “catch” items embedded in some of the questionnaires. A principal component analysis (PCA) was then performed on the psychometric data to reduce the dimensionality of the 11 questionnaire subscales: TCI_novelty-seeking_, TCI_harm-avoidance_, TCI_persistence_, FOP_minor_, FOP_severe_, FOP_medical_, PCS_rumination_, PCS_magnification_, PCS_helplessness_, STAI_state_, STAI_trait_. The data was centered and orthogonalized using a singular value decomposition algorithm. The influence of the 5 first components accounting for 92.5 % of the variance on pain valuation across the three experimental groups was then evaluated using multilevel regression as described above.

## Acknowledgments

Author contributions: H. Slimani, and M. Roy conceived the project. H. Slimani conducted the experiment with the help of Lana El Sabban. H. Slimani and M. Roy analyzed the data. H. Slimani, P. Rainville and M. Roy wrote the manuscript. All authors reviewed, edited, and approved the final version of the manuscript. This project was funded by the International Association for the Study of Pain, the Natural Sciences and Engineering Research Council of Canada, les Fonds de recherche du Québec: Nature et technologies and the Alan Edwards Centre for Research on Pain.

